# Modelling and analysis of cAMP-induced mixed-mode oscillations in cortical neurons: Critical roles of HCN and M-type potassium channels

**DOI:** 10.1101/2023.10.02.560623

**Authors:** Matteo Martin, Morten Gram Pedersen

## Abstract

Cyclic AMP controls neuronal ion channel activity. For example hyperpolarization-activated cyclic nucleotide–gated (HCN) and M-type K^+^ channels are activated by cAMP. These effects have been suggested to be involved in astrocyte control of neuronal activity, for example, by controlling the action potential firing frequency. In cortical neurons, cAMP can induce mixed-mode oscillations (MMOs) consisting of small-amplitude, subthreshold oscillations separating complete action potentials, which lowers the firing frequency greatly. We extend a model of neuronal activity by including HCN and M channels, and show that it can reproduce a series of experimental results under various conditions involving and inferring with cAMP-induced activation of HCN and M channels. In particular, we find that the model can exhibit MMOs as found experimentally, and argue that both HCN and M channels are crucial for reproducing these patterns. To understand how M and HCN channels contribute to produce MMOs, we exploit the fact that the model is a multiple-time scale dynamical system. We show that the MMO mechanism does not rely on the very slow dynamics of HCN and M channel gating variables, since the model is able to produce MMOs even when HCN and M channel activity is kept constant. In other words, the cAMP-induced increase in the average activity of HCN and M channels allows MMOs to be produced by the fast subsystem alone. We show that the fast-subsystem MMOs are due to a folded node singularity, a geometrical structure well known to be involved in the generation of MMOs in slow-fast systems. Besides raising new mathematical questions for multiple-timescale systems, our work is a starting point for future research on how cAMP signalling, for example resulting from interactions between neurons and glial cells, affects neuronal activity via HCN and M channels.

**Author summary:** Neurons use the frequency of electrical signals called action potentials to encode information, and various messenger systems interact with ion channels to control this so-called firing frequency. Recent experimental recordings show that the intracellular messenger cAMP can induce mixed-mode oscillations (MMOs) consisting of small-amplitude, subthreshold oscillations separating action potentials, which lowers the firing frequency greatly. We extend a recent mathematical model of neuronal electrical activity to investigate how MMOs occur from interactions between ion channels regulated by cAMP. Our simulations reproduce a range of experimental results, including cAMP-induced MMOs. We explain the model dynamics using modern geometrical methods that exploit the different timescales in the model. Our analyses show that the slow dynamics of cAMP-regulated HCN and M ion channels is not crucial for creating MMOs, but rather that the cAMP-induced increase in their average activity is important. Our analyses suggest that both HCN and M channels are crucial for MMOs and controlling the firing frequency, which has implications for our understanding of how astrocytes control neuronal information processing. Moreover, our study raises new mathematical questions related to how slow dynamical variables modify MMOs.

## Introduction

Cyclic AMP (cAMP) is an ubiquitous second messenger involved in a wide range of intracellular signaling processes. In neurons, cAMP has been suggested to control excitability and electrical activity by regulating ion channel activity [1, 2]. Of particular interest for the present work, in cortical neurons cAMP activates hyperpolarization-activated cyclic nucleotide–gated (HCN) channels [3, 4], which mediate a depolarizing current, and M-type potassium (also known as KCNQ or Kv7) channels, which carry a hyperpolarizing current [5, 6].

Recent experimental findings [4] demonstrated how Ca^2+^-evoked ATP release from astrocytes modulates the action potential (AP) conduction speed and the neuronal membrane excitability through cAMP increase and HCN channels. It was shown that HCN activation changes spiking electrical activity consisting of regular AP firing, into complex electrical patterns of mixed-mode oscillations (MMOs), where subthreshold, small amplitude oscillations (SAOs) intersperse large amplitude oscillations (LAOs), i.e., complete APs. This shift leads to a large reduction in the inter-spike frequency due to the presence of SAOs. Similarly, cAMP-induced activation of M-type channels has been shown to lower the firing frequency by producing MMOs [5]. How cAMP through both hyper- and depolarizing channels can cause MMOs is far from trivial.

This work proposes and investigates the hypothesis that the experimentally observed phenomena in [4, 5] result from both HCN and M channels. To study this idea, the neuronal electrical activity is modelled by the introduction of HCN and M currents in the model presented in [7]. This extended and optimized model replicates the experimental results presented in [4, 5]; in particular, it produces MMOs upon an increase in cAMP levels, in contrast to the simulations shown by Lezmy et al. [4].

To understand how M and HCN channels contribute to produce MMOs, we exploit that the model presents variables with different velocities, i.e., it is multiple-time scale dynamical system. In this class of models, geometrical and mathematical approaches are used to explain temporal dynamics, and the mechanisms generating MMOs are increasingly well understood [8]. Using these mathematical tools, it has been possible to understand and explain MMOs observed in many cellular quantities, e.g., complex patterns of electrical activity in neurons [9–11], pituitary cells [12–14], human beta cells [15], and cardiomyocytes [16, 17], as well as mixed-mode calcium oscillations [18], complex dynamics appearing from cell-to-cell interaction [19–21], etc.

We find that both HCN and M channels are important for creating MMOs in the model. However, this is not because of the dynamics of their gating variables, which operate on a very slow timescale, since the model is able to produce MMOs even when HCN and M channel activity is kept constant. In other words, cAMP increases the average HCN and M channel activity to set the fast subsystem in a region of parameter space where MMOs are produced by the fast subsystem alone. We show that the fast-subsystem MMOs are due to a folded node, and we construct numerically the relevant geometrical objects that explain the detailed dynamics of simulated MMOs.

## Results

### The model reproduces MMOs in various experimental conditions

We included HCN and M currents in a previous model of neuronal activity [7] in order to simulate and investigate electrical activity under different experimental conditions [4, 5], which examined how the cAMP-dependent pathway depicted in Fig 1 influence neuronal behavior. This model was also used in the study by Lezmy et al. [4] in which MMOs were observed experimentally in cortical neurons. Our simulations reproduce satisfactorily a range of biological results, and in particular exhibit MMOs as found experimentally.

**Fig 1.**
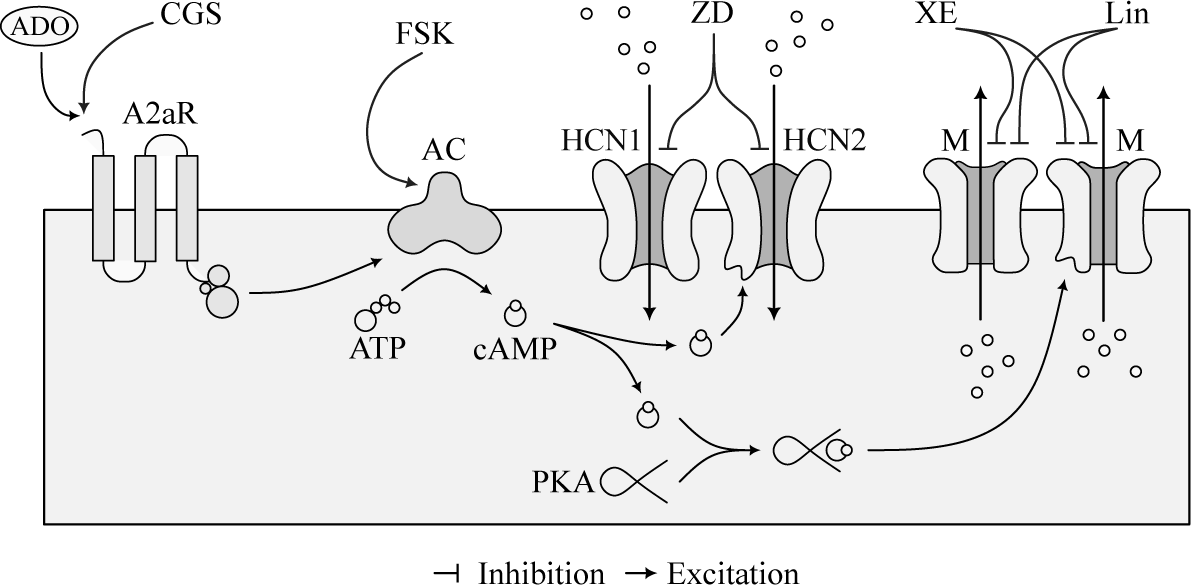
Schematic representation of the considered pathways. The scheme depicts how Adenosine 2A Receptor (A2aR) activation by extracellular Adenosine (ADO) increases cAMP levels via activation of adenylate cyclases (AC), which in turn increases the opening probability of HCN and M channels via direct binding and PKA activation, respectively. The experiments in [4, 5] target this pathway through specific drugs to understand how both M and HCN channels participate in the modulation of neuronal electrical activity. The chemical compounds employed are the A2aR agonist CGS21680 (CGS), the HCN channel inhibitor ZD7288 (ZD), the AC activator Forskolin (FSK), and the M channel antagonists XE991 (XE) and Linopirdine (Lin).

### Elevated cAMP following A2aR activation induces MMOs

In the first experiment, the control condition is compared to the experiment where a stimulus is provided via CGS21680 [4]. This drug activates A2aR and the downstream pathway, increasing the M and HCN currents, which can cause MMOs [4, 5].

The system of ODEs is solved for *I*_*App*_ equal to 250 and 300 *μ*A/cm^2^, and, in the presence of CGS, MMOs appear (Fig 2). The activation of the cAMP pathway augments the hyperpolarizing effect of the M current during the AP. In contrast, activation of HCN channels becomes important after the AP hyperpolarization phase. However, as will be shown in the following, although M and HCN currents are important for positioning the system so that MMOs can appear, it is not their dynamics that cause the SAOs. Rather, when the depolarizing HCN currents destabilize the resting membrane potential (RMP), the slow K^+^ current *I*_*KS*_ counteracts the depolarization, which in turn releases inactivation of the fast Na^+^ current *I*_*NaF*_ . The interplay between *I*_*KS*_ and *I*_*NaF*_ causes subthreshold oscillations, until *V* crosses the AP firing threshold. This model behavior reproduces the experimental evidence in [4].

**Fig 2.**
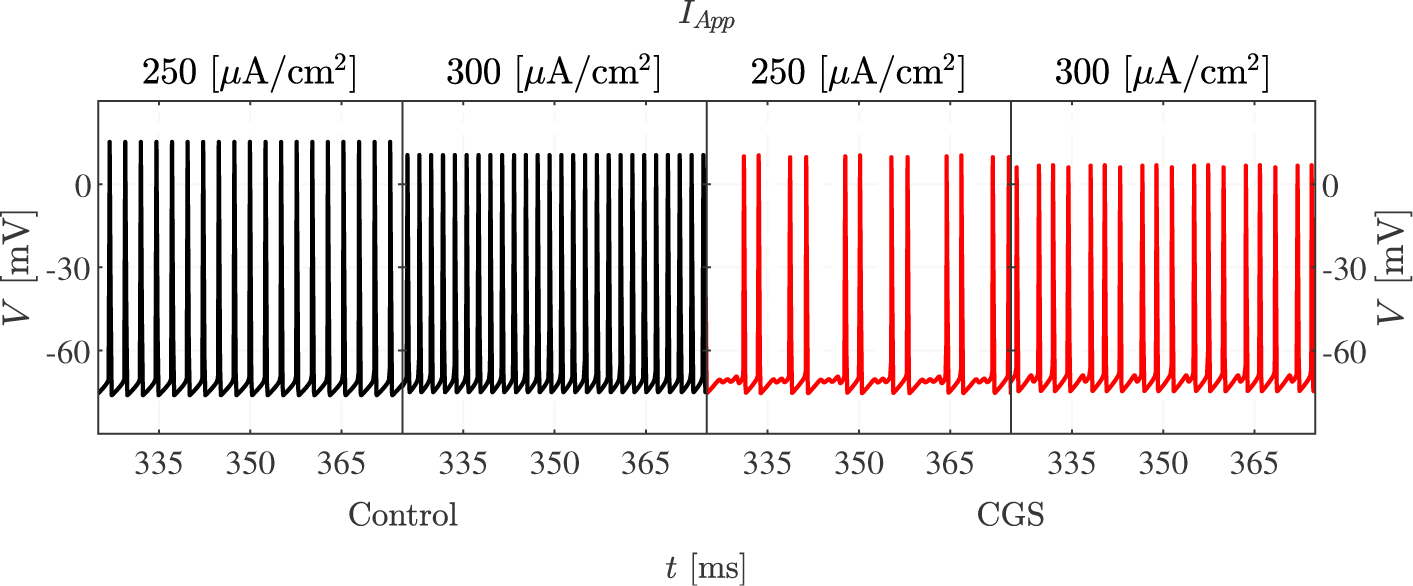
Elevated cAMP induces MMOs. The figure presents the simulated voltage traces under control conditions (left, black curves) and when cAMP is raised as in the experiments with CGS application (right, red) at two different values of *I*_*App*_.

### When HCN currents are blocked, cAMP can silence active cells

Lezmy et al. [4] tested how HCN currents influence neuronal responses by blocking them pharmacologically with ZD7288 under control conditions, or when the cAMP pathway was activated by CGS21680. In the model, this corresponds to increasing the M channel conductance only. Fig 3 presents simulated electrical activity under control and stimulated conditions. For *I*_*App*_ equal to 115 *μ*A/cm^2^, the control condition activates MMOs, whereas the addition of CGS21680 turns the neuron silent because of the activation of the hyperpolarizing M current. This simulation corresponds to the observation that in the presence of ZD7288 and at low stimulus strength, A2aR-mediated signalling from astrocytes reduced the frequency and even stopped neuronal AP firing [4].

**Fig 3.**
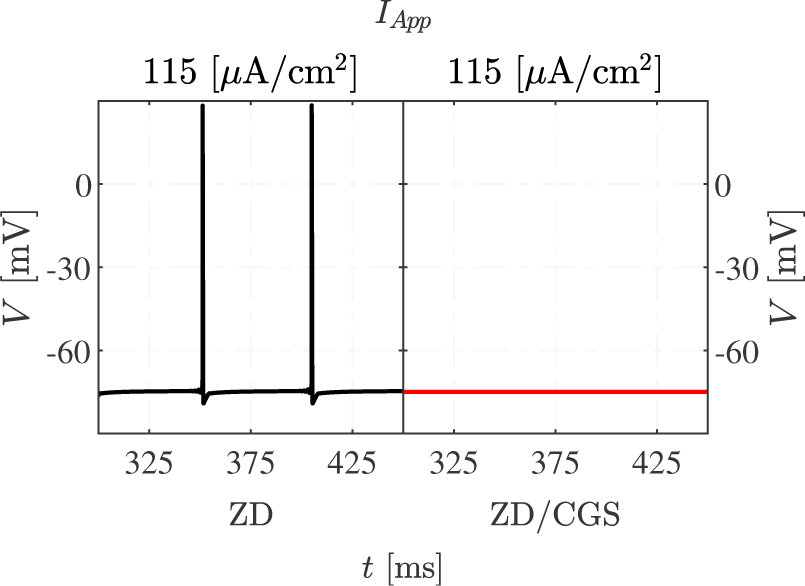
cAMP can stop AP firing when HCN are blocked. The figure presents the model simulations at relatively low *I*_*App*_ strength with HCN channels blocked in control (black) and CGS-stimulated (red) conditions. In this latter simulation, only M currents are activated.

### Blocking M channels changes MMOs to spiking and increases the firing frequency

Arnsten et al. [5] blocked M channels with XE991 to increase the comprehension of the roles of M and HCN channels. In the control state, no drug was applied to the neuron. We simulated this experiment by decreasing the M channel conductance, which increased the firing frequency greatly and lowered the AP amplitude (Fig 4). M-channel inhibition allows the HCN channel to depolarize the membrane potential quickly after the AP hyperpolarization phase. The fast Na^+^ channels do therefore not reactivate completely, which limits the following AP upstroke. Overall, the results in Fig 4 correspond to the experimental findings obtained in [5].

**Fig 4.**
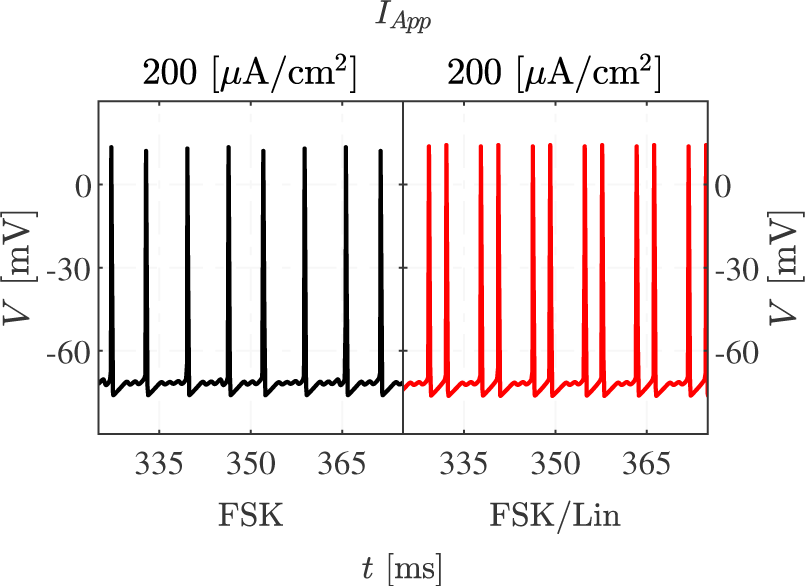
Blocking M channels increases the firing frequency. The figure presents model simulations in control conditions (left, black) and after inhibition of M channels (right, red), as in experiments with XE991 [5]. Depending on the size of *I*_*App*_, the increased firing frequency is caused by changing MMOs to spiking (*I*_*App*_ = 120 *μ*A/cm^2^), or by an acceleration of the rate of spiking accompanied by a slight reduction in AP amplitude (*I*_*App*_ = 320 *μ*A/cm^2^).

### Inhibiting M channels in stimulated conditions can change the signature of MMOs

To understand how M channels influence the observed dynamics with elevated cAMP, Forskolin, which raises the cAMP concentration, and the M-channel inhibitor Linopirdine were applied to the neurons [5].

Fig 5 presents the model results after the administration of Forskolin alone, or in the presence of both Forskolin and Linopirdine. Forskolin increases both *g*_*M*_ and *g*_*HCN*_, while Linopirdine decreases *g*_*M*_ . The partial inactivation of the M channel destabilizes the resting membrane potential and increases the excitability of the neuron. This effect explains the change in the MMO signatures in the presence of Linopirdine, with more LAOs and shorter sequences of SAOs compared to the scenario with fully active M channels.

**Fig 5.**
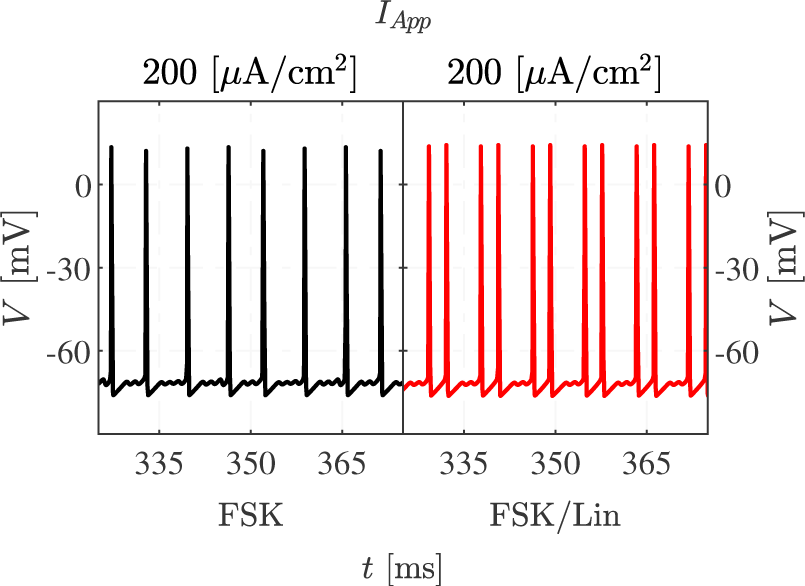
M-channel inhibition changes the signature of MMOs in stimulated conditions. Simulations of the model with raised cAMP in the absence (left, black) or presence of M channel inhibition, as in the experiments with Forskolin application in the absence or presence of Linopirdine [5], are presented. Note how the red trace presents more full APs (LAOs) and fewer subthreshold oscillations (SAOs).

### Firing frequency analyses

Fig 6 shows firing frequency (FF) curves under different experimental conditions for a range of *I*_*App*_ values. The FF curve associated with the unstimulated neuron (black) splits into several parts. For *I*_*App*_ in the interval [81, 224] *μ*A/cm^2^, the neuron exhibits MMOs. The neuron is silent if *I*_*App*_ is below 81 *μ*A/cm^2^, whereas for *I*_*App*_ ∈ [224, 720] *μ*A/cm^2^, only LAOs persist, i.e., the cell produces regular spiking. The application of CGS21680 activates A2aR, which raises cAMP levels, increasing both HCN and M currents. This modification changes the FF curve (red) by shifting it leftward and making it less steep. In fact, the unstimulated FF curve crosses the one obtained for CGS21680 stimulation at *I*_*App*_ ≈ 113 *μ*A/cm^2^. Above this value the unstimulated case presents higher FF, while the stimulated one is more active for *I*_*App*_ below the threshold. This crossing of the FF curves agrees with the results by Lezmy et al. [4].

**Fig 6.**
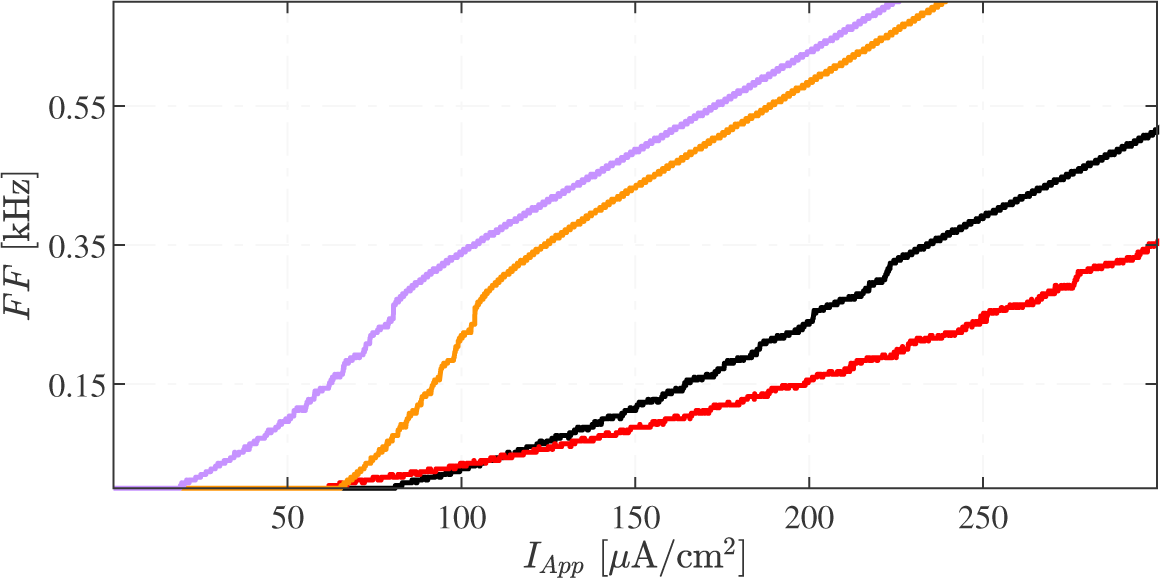
Firing Frequency analyses. The FF of the 5D model under various pharmacological stimulations at different levels of applied current. The black curve shows FFs for the unstimulated (control) condition. Red, orange and violet curves are associated with GCS21680 (increased cAMP), XE991 (M channel block), and simultaneous XE991+CGS21680 application, respectively.

Fig 6 shows the FF results for two additional conditions. M currents are inhibited in both these cases, mimicking application of XE991 in the absence (orange) or presence (violet) of the A2aR agonist CGS21680. Considering the XE991-treated neuron, the associated FF curve is steeper compared to the control and the CGS21680-treated cases. In fact, under M-channel inhibition, the neuron generates oscillatory phenomena for lower *I*_*App*_, due to the lack of the stabilizing, hyperpolarizing M current. For *I*_*App*_ in [65, 80] *μ*A/cm^2^, the unstimulated neuron is silent, while the one treated with XE991 undergoes MMOs with a FF approximately proportional to *I*_*App*_. If the neuron pre-treated with XE991 undergoes CGS21680 stimulation, the FF curve shifts further to the left. The increase in the depolarizing HCN channel conductance explains this movement, since the neuron is more excitable. The comparison between the curves presented in Fig 6 and the experimental FF curves [4, 5], strongly suggests that both M and HCN currents are required to explain how cAMP controls the FF in cortical neurons.

### Dynamical system analyses of the fast subsystem

To understand how MMOs appear in the 5D model simulated above, we analyse the three-dimensional fast (*V, h, s*) subsystem of the 5D model, which allows us to apply standard methods regarding folded singularities [8]. Exploiting the fact that the M and HCN channels have slower dynamics than the other variables in the model, the activation variables of M (*w*) and HCN (*r*) channels are considered as parameters in the fast (*V, h, s*) subsystem (the “3D model” in the following). The reduction steps are illustrated in the section Model Reduction in Materials and Methods. Figure 7 presents the 3D model simulation of the experiment with CGS stimulation, corresponding to the 5D model simulation shown in Fig 2. Crucially, the dynamics of HCN and M channels is not strictly necessary in order to observe MMOs, which is a result of the dynamics of the 3D fast subsystem. The most significant difference between the 5D (Fig 2) and the 3D (Fig 7) simulations are seen in the signatures of the MMOs presented by the two systems. Specifically, for the same value of *I*_*App*_, the 3D model shows less LAOs and more SAOs than the 5D model. This variation is caused by the simplification of the model.

**Fig 7.**
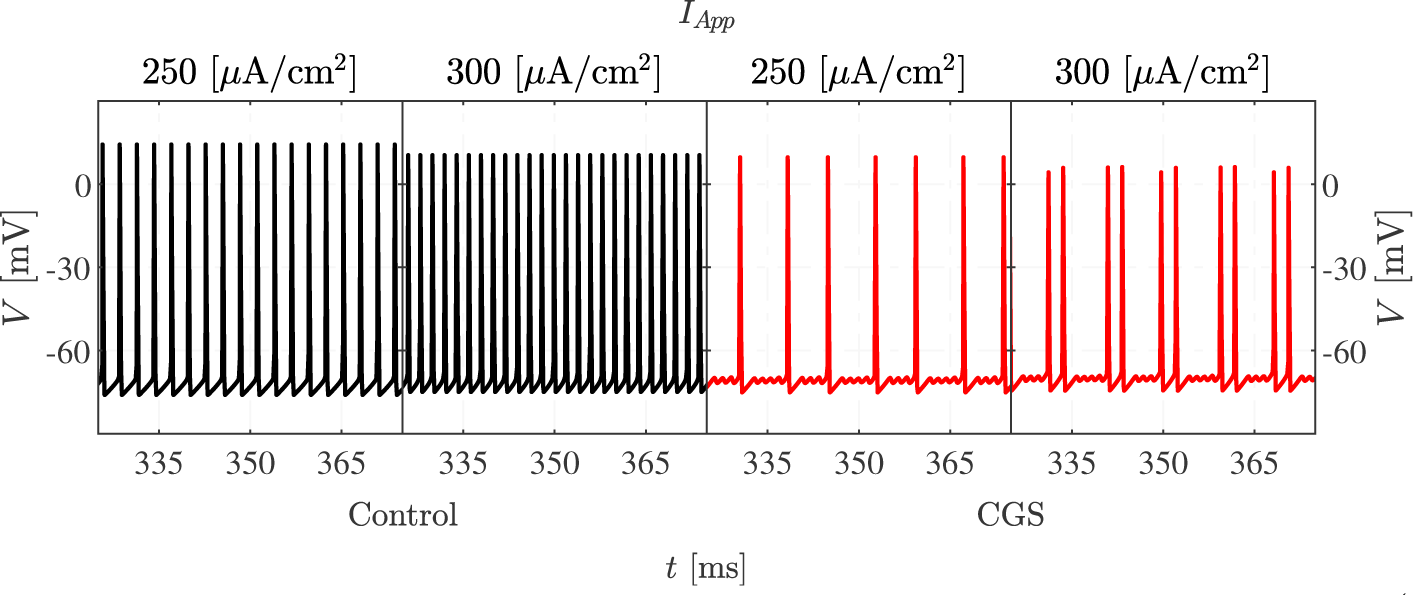
3D model voltage traces. Simulations of the 3D model under control (left, black) and CGS-stimulated (right, red) conditions, compare with Fig 2.

In order to be able to use the same fast-subsystem bifurcation diagrams in the presence and absence of cAMP-mediated activation of M and/or HCN channels, we introduce the modified slow variables (where *cAMP* = 0, respectively *cAMP* = 1, indicate absence, respectively presence, of cAMP-mediated channel activation)

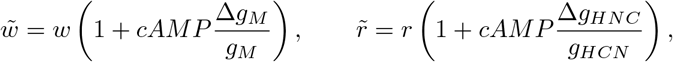

which are then used as bifurcation parameters in the fast subsystem.

### Fast-subsystem bifurcation diagrams

When projecting the trajectory of the 5D model onto the 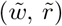) plane, the model trajectory evolves close to a straight line 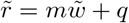 (Fig 8A). We can therefore construct one-parameter bifurcation diagrams (1P-BD) for the 3D fast subsystem with bifurcation parameter 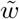, and 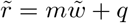 constrained to lie on the identified straight line. However, the line depends on the experimental condition that is simulated, and moves upwards when cAMP is elevated (Fig 8A): Both 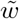 and 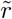increase since cAMP introduces a shift in these variables by construction.

**Fig 8.**
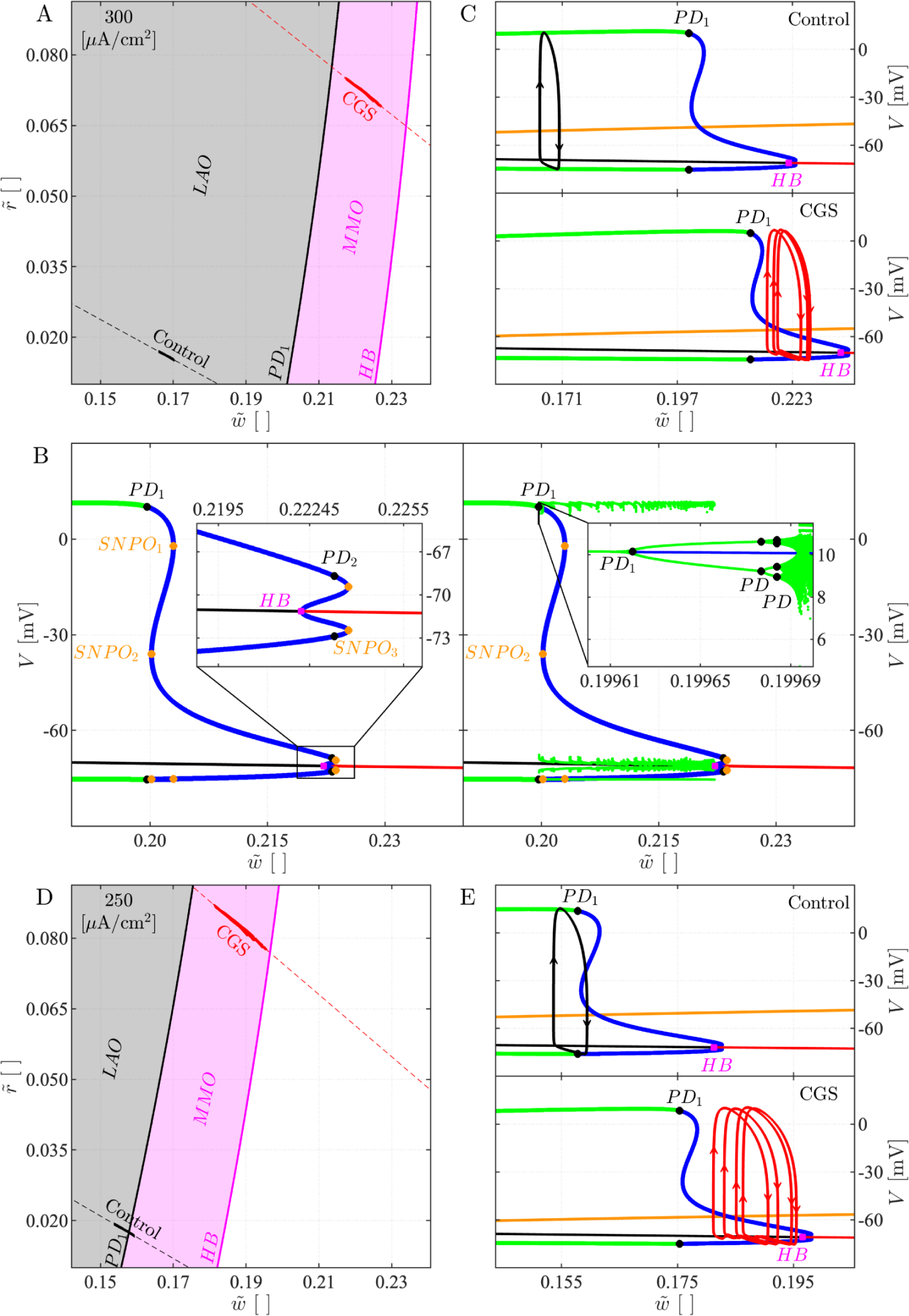
Fast 3D subsystem bifurcation diagrams. A: 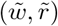 plane with projections of 5D model simulations for *I*_*App*_ = 300 *μ*A/cm^2^ in control (full, black curve) and CGS-stimulated (red) conditions. The dashed lines indicate the linear approximations 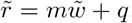. The background shows the 2-parameter bifurcation diagram (2P-BD) for the fast subsystem with 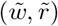 as parameters, obtained by following the most relevant bifurcations found in the 1P-BD (panel B). (Caption continues on next page).

Fig 8B shows the computed 1P-BD for the fast (*V, h, s*) subsystem. The results show the existence of a unique equilibrium point. For high M-channel activation 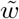, the equilibrium is stable. As 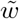 is reduced (and 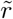 is increased), the equilibrium loses stability in a Hopf bifurcation (HB). From the HB, unstable periodic solutions emerge. On the other hand, at very low 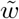, the branch of unstable equilibria is surrounded by a branch of stable limit cycles, and as 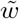 is increased, these lose stability as they go through a cascade of period doubling bifurcations (PDs). Following saddle-node of periodic orbit (SNPO) bifurcations, the branch of (unstable) limit cycles eventually connects to the HB point. Between PD_1_ and HB, the system exhibits MMOs. These do not appear as a direct result of the PDs, but rather when the chaotic attractor makes a sudden discontinuous jump slightly to the right of PD_1_ at 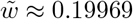.

Reintroducing the slow dynamics of 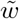 (and 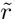), we see that for *I*_*App*_ = 300 *μ*A/cm^2^ in the control condition, the system follows a stable periodic orbit without exhibiting SAOs (Fig 8C, upper). In the case with CGS stimulation, the simulated trajectory is fully contained within the (3D fast-subsystem) MMO region (Fig 8C, lower): 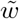 decreases during the SAOs, but increases during the LAOs (APs), so that 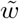 remains in the interval with MMOs. Geometrically, the explanation is that in the presence of cAMP (*cAMP* = 1), the 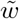 nullcline is moved down and to the right (Fig 8C), so that the system stalls further to the right in the MMO region, rather than in the LAO region as with *cAMP* = 0.

The behavior with lower applied current (*I*_*App*_ = 250 *μ*A/cm^2^) is interesting. Here, the fast-subsystem bifurcations occur at lower values of 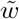. Due to the evolution of the dynamical variables 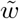 and 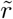, the 5D system enters the region with fast-subsystem MMOs (Fig 8D). However, the system does not stay long enough in this region for SAOs to appear, since 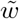 decreases for low *V* and the trajectory moves to the left of PD_1_ where an AP appears (Fig 8E, upper). With CGS stimulation (*cAMP* = 1; Fig 8E, lower), as in the case with *I*_*App*_ = 300 *μ*A/cm^2^, the trajectory lies completely in the MMO region since the 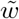 nullcline lies lower and to the right, compared to the control case (*cAMP* = 0).

Relaxing the constraint 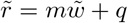 permits construction of the two-parameter 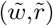 BD of the fast (*V, h, s*) subsystem by following the main bifurcations shown in the 1P-BD (Fig 8AD). MMOs are observed only if the entire 5D model trajectory belongs to the MMO region. The control condition with *I*_*App*_ = 250 *μ*A/cm^2^ is a borderline scenario. In fact, as discussed above, the trajectory crosses the PD_1_ curve but presents no SAOs (Fig 2). Altogether, the retrieved 2P-BD for the 3D fast subsystem reflects the full 5D model behavior well, and suggest that the average activation level of HCN and M channels is sufficient to predict the type of activity. For example, activation only of M channels corresponds to a right-shift in the 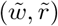 plane, which could move the system from the MMO region to the silent region to the right of the HB curve, as seen in Fig 3. Vice versa, blocking M channels corresponds to a left-shift, which can cause the system to go from the MMO to the LAO region, as seen in Fig 4. Moreover, activation solely of the HCN current corresponds to an increase in 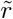 only, i.e., an upwards step in the 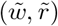 plane, which could activate a silent cell to exhibit MMOs, or change MMOs to simple spiking by moving the system into the LAO region. This explains the results shown in Fig 6 for XE991 versus XE991+CGS21680.

The derived BDs can be interpreted biophysically as follows. As the depolarizing HCN channels activate (higher 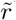), and the M channels become less active (lower 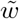), the cell’s excitability increases, facilitating AP generation. This observation explains why stable periodic orbits occur at a low 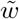 and high 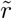. Instead, when 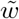 increases and 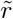 decreases, the excitability of the model reduces. In addition, the more pronounced M channel activation is, the more SAOs appear, until the equilibrium corresponding to the resting potential eventually becomes stable for high 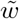values.

**Fig 8**. (continued) *B* : 1P-BD of the fast subsystem for *I*_*App*_ = 300 *μ*A/cm^2^ in the control condition with 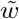 as parameter and 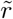 constrained to the dashed, black line in panel A. The red (respectively, black) curve indicates stable (unstable) equilibria, the green (blue) curves are minima and maxima of stable (unstable) periodic solutions. The inset in the left panel shows a zoom on the region where the equilibrium changes stability. In the right panel, local maxima and minima of 3D model simulations are added. The inset shows a zoom on the period-doubling cascade. HB: Hopf bifurcation, SNPO: saddle-node bifurcation of periodic orbits, PD: period doubling bifurcation. *C* : BD as in panel B with 5D simulation projected onto the 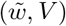 plane for the control (upper) and CGS-stimulated (lower) scenarios. The orange curve is the 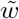 nullcline. *D* and *E* : As panels A and C, but for *I*_*App*_ = 250 *μ*A/cm^2^.

### M currents are necessary to reproduce the experiments

The model parameters are taken from [4, 7, 22]. However, the HCN reversal potential, *E*_*HCN*_, the M-current conductance, *g*_*M*_, and the increase in this conductance due to cAMP, ∆*g*_*M*_, are fine-tuned to satisfy both physiological and mathematical constraints to reproduce the electrical activity observed in the experimental study of [4] (Fig 2). The physiological restrictions require *E*_*HCN*_ to be within the range [−50, −20] mV [23], while the cAMP-induced increase in M-current conductance is less than the conductance in control conditions, i.e., ∆*g*_*M*_ ≤ *g*_*M*_ [5, 24]. *The mathematical constraints should guarantee the reconstruction of the experimentally observed voltage traces [4]*. *That is, we look for a combination of (E*_*HCN*_, *g*_*M*_, ∆*g*_*M*_) that generates spiking in control condition but MMOs when cAMP is raised, for example after CGS application [4].

By fixing *E*_*HCN*_ and *g*_*M*_, the modification of ∆*g*_*M*_ only affects the dynamics in the stimulated condition. As seen in the 2P-BD with bifurcation parameters *I*_*App*_ and ∆*g*_*M*_ with *g*_*M*_ = 50 mS/cm^2^ and *E*_*HCN*_ = − 50 mV (Fig 9), higher values of ∆*g*_*M*_ expand and right-shift the interval where the model activates MMO at elevated cAMP. Similar results were found for other values of *g*_*M*_ and *E*_*HCN*_ . In order for the model to produce, at some *I*_*App*_, MMOs at elevated cAMP but spiking in control conditions, the area with MMOs at high cAMP must go further to the right than the MMO area for the control case. In Fig 9 this happens only for ∆*g*_*M*_ *>* 11 mS/cm^2^, and as ∆*g*_*M*_ increases the interval of *I*_*App*_ values where spiking is seen in control condition but MMOs in stimulated condition widens. We used ∆*g*_*M*_ = *g*_*M*_ = 50 mS/cm^2^.

**Fig 9.**
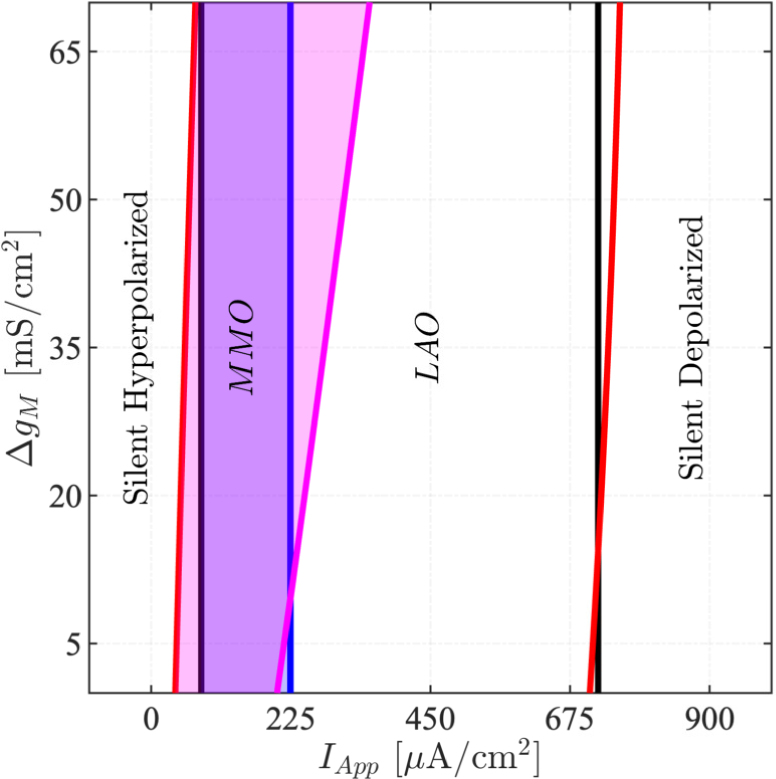
Model dynamics depends on the degree of cAMP-activation of M channels. Two-parameter BD with bifurcation parameters (*I*_*App*_, ∆*g*_*M*_) for *g*_*M*_ = 50 mS/cm^2^ and *E*_*HCN*_ = − 50 mV. Black and red curves indicate respectively HBs in the control conditions and with raised cAMP. The curves in blue and magenta represent, similarly, the PDs (PD_1_ in Figs 8) in the control and stimulated cases. The combination of *I*_*App*_ and ∆*g*_*M*_ where MMOs occur is highlighted with shaded blue (control) and magenta (elevated cAMP) areas.

Summarizing, if cAMP does not affect the M current but HCN channels only, the model is not able to reproduce the experimental finding that CGS via cAMP elevation switches spiking electrical activity to MMOs.

### Geometry of mixed-mode oscillations

The 5D model is a multiple time-scale dynamical system as noted above. Its 3D fast subsystem is itself a slow-fast system that presents one fast (*V*) and two slow (*h* and *s*) variables. We now explain the local and global dynamics of this 3D system with particular attention to the origin of MMOs. For a brief introduction to the underlying theory see Materials and Methods, and, for in-depth expositions, refs. [8, 25].

Among the model’s geometrical structures, the critical manifold 𝒞_0_ is of great importance in order to understand the dynamics of the system. It is defined as the set of points (*V, h, s*) where *dV/dt* = 0, i.e., the set of equilibrium points for the fast subsystem of the 3D model. Points on 𝒞_0_ are said to be attracting or repelling if they are so when interpreted as fast-subsystem equilibrium points. For our model, the critical manifold can be expressed explicitly as a graph,

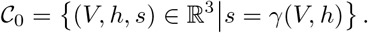

We find, in the physiologically relevant region, that *𝒞*_0_ has a folded structure with two folds, *ℒ* ^−*/*+^. These folds split *𝒞*_0_ into attracting and repelling sheets, denoted respectively 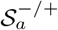 and 𝒮_*r*_. Starting from *V* close to the neuron’s resting potential, 𝒞_0_ is thus decomposed as 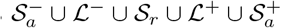.

At the fold, the system becomes singular (see Materials and Methods and [8]). To understand the dynamics near the fold, the system is therefore desingularized, and it turns out that equilibrium points of the desingularized system on a fold, so-called folded singularities, play a crucial role in understanding the origin of SAOs [8]. We find that the model possesses a folded node (FN) singularity, which is well-known to cause *canard-induced* SAOs [8].

Let 0 *< ϵ* ≪ 1 denote the time scale separation between the fast *V* variable and the slower *h* and *s* variables, see also Materials and Methods. The following analyses assume that the model’s initial conditions are located near 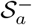. Assuming stability of the 3D model equilibrium, i.e., for all 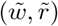 located on the right of the HB curve in Fig 8, the 3D fast-subsystem trajectory evolves dominated by the slow flow constrained to an 𝒪 (*ϵ*) perturbation of the attracting submanifold 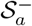, denoted 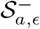, and approaches the stable equilibrium in infinite time.

Instead, if the equilibrium point is unstable, the dynamics depends on how the system approaches the fold ℒ ^−^. For parameter combinations in the spiking region, e.g., the control condition shown in Fig 7 and Fig 10 (upper panels), the system’s trajectory evolves constrained to 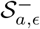 until it reaches the fold curve *ℒ* ^−^ at a *regular jump point* [8, 25], *where it then switches dynamics and moves to* 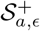 via its fast flow. In Fig 10, this corresponds approximately to crossing *ℒ* ^−^ to the left of the FN. On 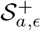 it again evolves according to the slow flow. When it reaches ℒ ^+^, the system jumps to 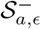, and then the above steps repeat, creating relaxation oscillations.

**Fig 10.**
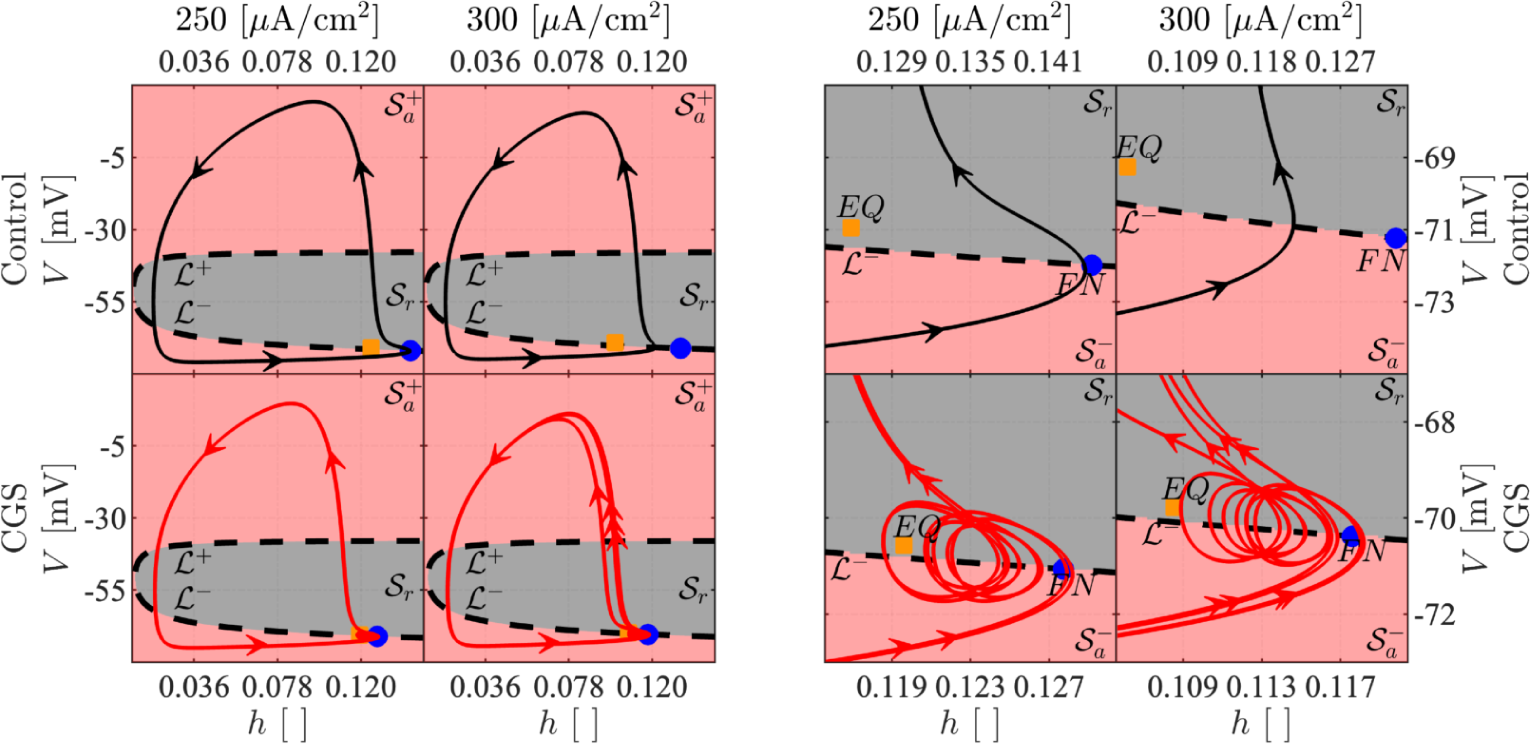
Critical manifold and folded singularities of the 3D model. *The presented phase-plane plots show the trajectory of the 3D model with parameters as in Fig 7 projected onto the (h, V*) plane for the control case (black curve, upper panels) and in the presence of CGS (red, lower), observed globally (left) and locally with a zoom on the region where SAOs appear for the CGS-stimulated case (right) The dashed black curve represents the fold *ℒ* ^−*/*+^ of the critical manifold. The red and gray shaded regions correspond to the attracting, _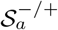_, and repelling, 𝒮 _*r*_, submanifolds, respectively. The unstable equilibrium point is given by the orange square, while the blue dot indicates the FN. The arrows indicate the direction of the flow.

If on the otherhand the system approaches *ℒ* ^−^ near the folded node, more precisely in the *funnel region*, SAOs are produced as the system passes from 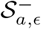 to *S*_*r,ϵ*_ [8, 15, *25]*.

The funnel is divided into *rotational sectors* bounded by canards, and the number of SAOs depends on the sector in which the folded node is approached. The trajectory eventually jumps to 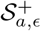, corresponding to the onset of an AP, i.e., a LAO, and then follows the slow flow until it reaches ℒ ^+^ and jumps back to 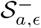. The trajectory may reenter the funnel region, producing single LAOs separated by SAOs, or it may reach ℒ ^−^ at a regular jump point, thus producing two or more LAOs separated by SAOs, as in Figs 7 and 10 with CGS for *I*_*App*_ = 250 and *I*_*App*_ = 300 *μ*A/cm^2^, respectively.

In more details, the dynamics shown in Figs 7 and 10 can be understood as follows. Each rotational sector *R*_*i*_ is bounded by two canard solutions, *ξ*_*i*−1_ and *ξ*_*i*_, where *i* indicates the number of SAOs exhibited by the canard. In Fig 7, the system at *I*_*App*_ equal to 250 and 300 *μ*A/cm^2^ evolves with *signatures* 1^4^1^3^ and 2^4^2^3^, respectively. In the former, the model generates 1 LAO followed by 3 or 4 SAOs. Thus, the return mechanism projects alternatingly the trajectory into *R*_4_, delimited by *ξ*_3_ and *ξ*_4_, and into *R*_3_, delimited by *ξ*_2_ and *ξ*_3_, as presented in Fig 11. Instead, in the case of *I*_*App*_ = 300 *μ*A/cm^2^, 2 LAOs occurs, followed by 3 or 4 SAOs. The generation of 2 consecutive LAO is related to the return mechanism. Indeed, after the first large excursion, the system is projected to the left of *ξ*_0_, which bounds the FN funnel. This fact implies that the trajectory evolves onto 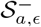 until it meets *ℒ* ^−^ at a regular jump point, from where it jumps to 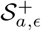 without making SAOs. After the second LAO, the trajectory is projected into the FN funnel either and alternatingly between *ξ*_2_ and *ξ*_3_ (*R*_3_), or between *ξ*_3_ and *ξ*_4_ (*R*_4_), where 3, respectively 4, SAOs occur.

**Fig 11.**
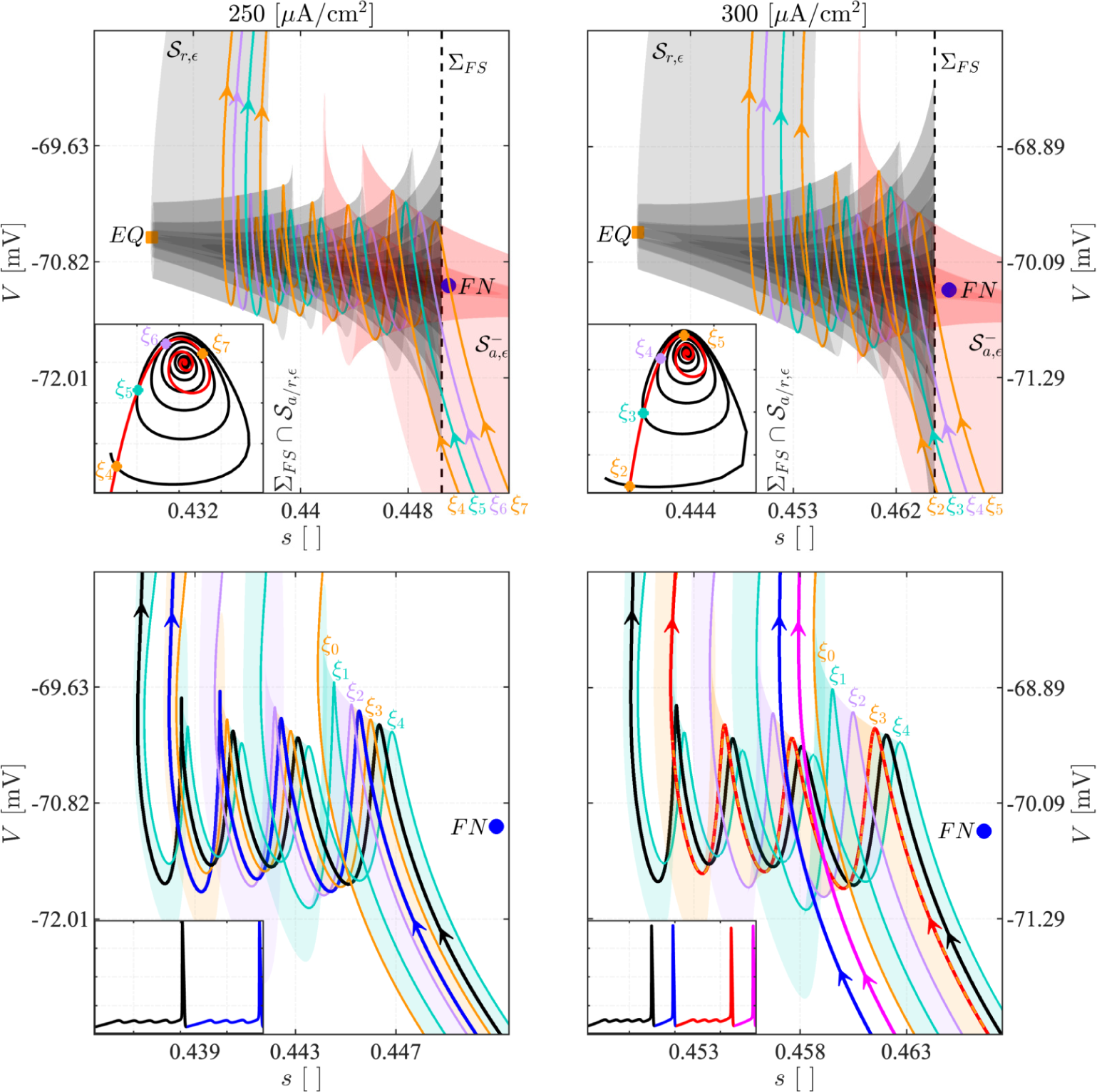
Canards and attracting and repelling manifolds for the 3D Model. The upper panels show the local reconstruction of 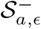 (shaded red) and _*r,ϵ*_ (shaded black) for *I*_*App*_ = 250 *μ*A/cm^2^ (left) and *I*_*App*_ = 300 *μ*A/cm^2^ (right). In each panel, the blue dot indicates the FN. The vertical black dashed line identifies a plane Σ_*F S*_ near the FN, and the inset presents the corresponding intersections between Σ_*F S*_ and, respectively, 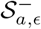 (red) and 𝒮 _*r,ϵ*_ (black). The colored lines *ξ*_4_ − *ξ*_7_ and *ξ*_2_ − *ξ*_5_, obtained for *I*_*App*_ equal to 250 and 300 *μ*A/cm^2^, respectively, represent the canard orbits identified via the intersection points between 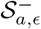 and *𝒮* _*r,ϵ*_ in the plane Σ_*F S*_. In the lower panels, the canard orbits *ξ*_0_ − *ξ*_4_, the associated rotational sectors *R*_1_ − *R*_4_ and the local dynamics of the 3D model are presented for the considered values of *I*_*App*_. The trajectory’s temporal and phase-plane evolutions are color-coded to highlight how the different signatures relate to the identified geometrical structures.

## Discussion

In this study we developed a model of electrical behavior of a neuronal cell subjected to different treatments interacting with the cAMP-dependent pathway linking A2aR signalling to HCN and M channel activation. The model parameters were chosen in compliance with biological constraints to ensure that the model aligns with physiological knowledge. The model qualitatively reproduced the experimental results [4, 5], validating its utility in elucidating the role of HCN and M channels.

We found that both channels shape the neuronal electrical activity. With enhanced HCN activation, the neuron’s excitability increases, thereby facilitating the generation of APs. When the M current predominates, the resting membrane potential becomes more stable. The activation of these two currents can give rise to complex electrical phenomena such as MMOs. Our analyses strongly support the idea that both HCN and M channels are needed for producing MMOs corresponding to experimental results. Even if these temporal patterns are fragile, they are crucial for controlling the neuronal firing frequency. We thus suggest that when neuronal and glial cells interact [4], both these two channels are involved to allow glial cells to finely tune the electrical behavior in individual neurons.

To explain the model behavior, we exploited the fact that we were dealing with a multiple timescale system. The 3D fast subsystem of the 5D model, obtained by fixing HCN and M channel gating variables to their average values, was shown to be able to produce MMOs due to the presence of a folded node . Hence, it is not the slow dynamics of HCN and M channel activity that cause MMOs in the model, but rather, cAMP increases the average activity of these channels, setting the 3D fast subsystem in a region where MMOs occur. We showed how the bifurcation diagrams for the 3D fast subsystem with the modified gating variables, 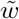 and 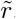, provided insight into various experiments that were reproduced with simulations of the full 5D model.

The 3D model qualitatively reproduced the behavior of the 5D system as presented in Fig 7, despite some discrepancies in the generated MMO signatures. We speculate that these differences are related to the assumption of constant HCN and M channel gating variables, *r* and *w*. Reintroducing the dynamics of these variables slowly modulates the location and the properties of the folded node and its funnel, which would modify the MMO signature. For a mathematical study of this idea, numerical continuation methods may be valuable tools to continue the canard orbits of the 3D reduced model by varying the parameters 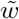 and 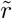. Such continuation would provide insights regarding how the FN funnel changes as the model parameters are changed. For this scope, it may be convenient to use a 4D model, such as the one obtained by constraining one of the HCN and M channel activation variables to a straight line, whilst the other is free to evolve with its dynamic. This framework could apply geometrical theories and bifurcation analyses similar to the ones used here to understand the local and global evolution of the model.

From a biological point of view, it would be interesting to investigate theoretically what happens if cAMP levels depend on dynamics of glial cells, and how the different types of single-cell electrical phenomena impact the overall behavior of small and large networks.

In conclusion, this study employed a white-box realistic neuronal cell model to unveil the role of HCN and M channels in shaping electrical behavior as a result of cAMP signalling. We demonstrated and analyzed how the resulting currents can influence and evoke phenomena such as MMOs. This work represents a starting point for future research on the interplay between these two channels and their regulation of neuronal electrical activity, as well as for investigations on how cAMP signalling, for example resulting from interactions between neurons and glial cells, affects neuronal activity.

## Materials and Methods

### Model

The core structure of the model is presented in Richardson et al. [7]. However, the activation variables of both fast and persistent Na^+^ channels are simplified with their steady-state functions, as shown below. The model equations are

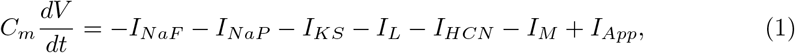

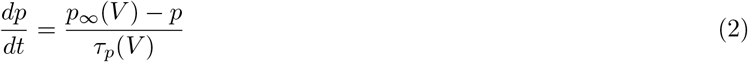

where *V* refers to the neuron’s membrane potential, *t* is the time variable, *C*_*m*_ is the membrane capacitance, and *p* ∈ *{h, s, r, w}* indicates a general activation/inactivation variable. The right-hand side of the first ODE is a sum of ionic currents. These quantify respectively the fast and persistent Na^+^ currents (*I*_*NaF*_ and *I*_*NaP*_), the slow K^+^ current (*I*_*KS*_), the leakage current (*I*_*L*_), the HCN current (*I*_*HCN*_), the M-type potassium current (*I*_*M*_) and finally the applied current (*I*_*App*_). These currents follow

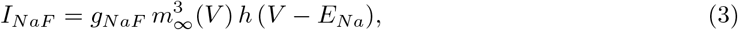

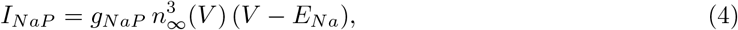

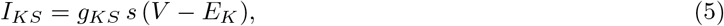

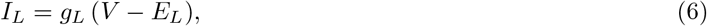

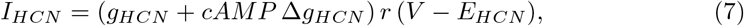

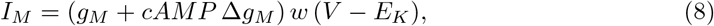

with *m*_∞_, *n*_∞_, *s, r, w* activation variables of fast Na^+^, persistent Na^+^, slow K^+^, HCN and M channel currents, whereas *h* is the *I*_*NaF*_ inactivation variable. The terms *E*_*X*_ with *X* ∈ *{Na, K, L, HCN}* represent the Nernst potential for sodium, potassium, leakage and HCN currents, respectively, and *g*_*X*_ with *X* ∈ *{NaF, NaP, KS, L, HCN, M}*, indicate the maximal whole-cell conductances.

∆*g*_*HCN*_ and ∆*g*_*M*_ model, respectively, the alteration of HCN and M maximal conductances after a rise of the intracellular cAMP levels, as indicated by the binary variable *cAMP* . Finally, the terms *p*_∞_ and *τ*_*p*_ refer to the activation/inactivation steady-state functions and time constants, respectively. For each *p* ∈ *{h, s, r}*,

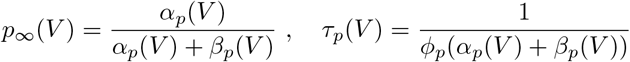

where *α*_*p*_ and *β*_*p*_ represent the transition rates of the activation/inactivation variable, and *ϕ*_*p*_ indicates the temperature correction factor which obeys the following law [26],

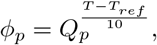

where *Q*_*p*_ and *T*_*ref*_ are channel-dependent terms, and *T* is the characteristic temperature of the experiment. The *α*_*p*_ and *β*_*p*_ are given by

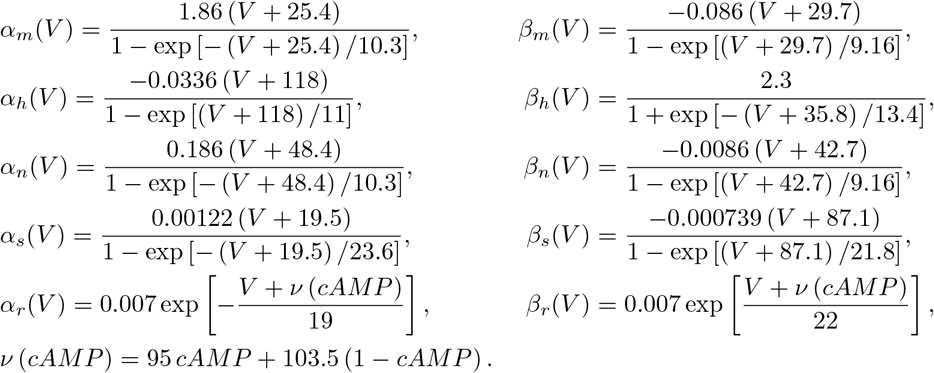

Finally, the steady state (*w*_∞_) and the time constant (*τ*_*w*_) of the *w* activation variable are [22]

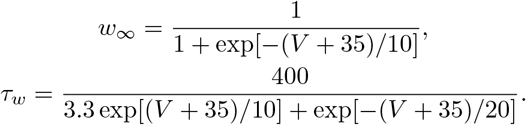

Table 1 presents the model parameters, obtained from [4, 7, 22]. To simulate the different experimental scenarios, the parameters presented in Table 2 were used. The ODE system was solved numerically with MATLAB [27] and XPPAUT [28]. Specifically, XPPAUT was used to derive the bifurcation diagrams presented in Figs 8 and 9, using a fourth-order Runge-Kutta method with a time step equal to 0.01 ms. MATLAB executed successive data post-processing, visualization and slow-manifold reconstruction algorithms. The system of ODEs was solved with the built-in MATLAB functions ode45 and ode78.

**Table 1.** Default model parameters. *T*_(*ref*,1)_ is the reference temperature for the activation/inactivation properties of the slow-K^+^ and fast-Na^+^ channels, while *T*_(*ref*,2)_ is associated with the HCN channels.

| Parameter | Value | Unit of Measure | Parameter | Value | Unit of Measure |
| --- | --- | --- | --- | --- | --- |
| $g_{NaF}$ | 1000 | [mS/cm<sup>2</sup>] | $T$ | 37 | [°C] |
| $g_{NaP}$ | 1 | [mS/cm<sup>2</sup>] | $T_{(ref,1)}$ | 20 | [°C] |
| $g_{KS}$ | 40 | [mS/cm<sup>2</sup>] | $T_{(ref,2)}$ | 35 | [°C] |
| $g_L$ | 11.3 | [mS/cm<sup>2</sup>] | $Q_h$ | 2.9 | [ ] |
| $g_{HCN}$ | 23 | [mS/cm<sup>2</sup>] | $Q_s$ | 3.0 | [ ] |
| $g_M$ | 50 | [mS/cm<sup>2</sup>] | $Q_r$ | 3.0 | [ ] |
| $\Delta g_{HCN}$ | 12 | [mS/cm<sup>2</sup>] | $E_{Na}$ | 50 | [mV] |
| $\Delta g_M$ | 50 | [mS/cm<sup>2</sup>] | $E_K$ | -84 | [mV] |
| $C_m$ | 0.9 | [μF/cm<sup>2</sup>] | $E_L$ | -83.38 | [mV] |
| $I_{App}$ | | [μA/cm<sup>2</sup>] | $E_{HCN}$ | -50 | [mV] |

**Table 2.**
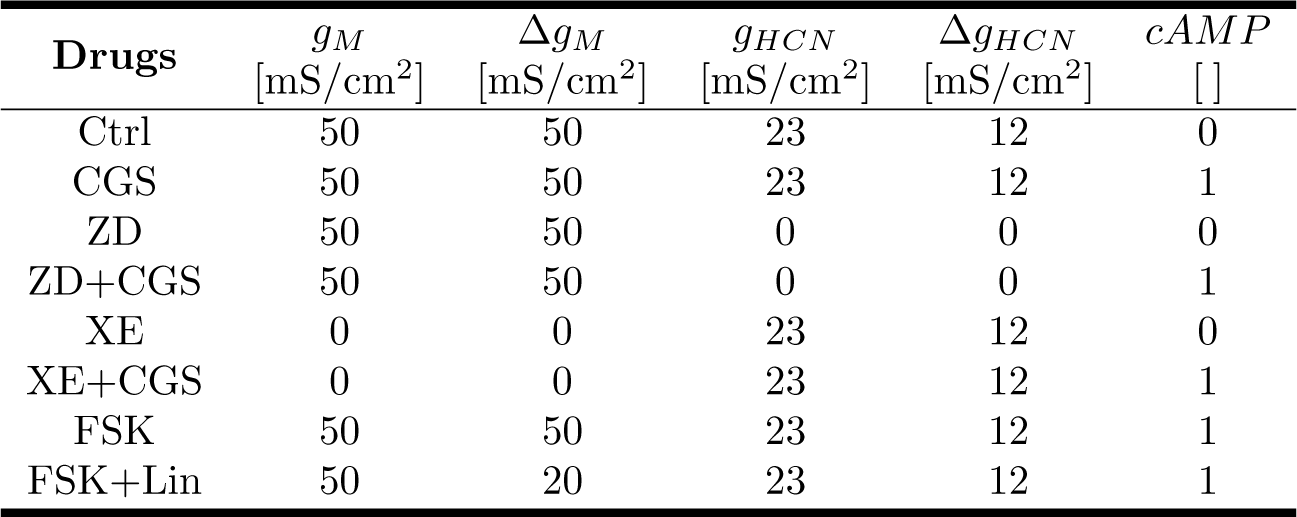
Translation of drug combinations applied in the various experiments into parameter sets used to simulate the model.

| Drugs | $g_M$<br>[mS/cm<sup>2</sup>] | $\Delta g_M$<br>[mS/cm<sup>2</sup>] | $g_{HCN}$<br>[mS/cm<sup>2</sup>] | $\Delta g_{HCN}$<br>[mS/cm<sup>2</sup>] | $cAMP$<br>[ ] |
| --- | --- | --- | --- | --- | --- |
| Ctrl | 50 | 50 | 23 | 12 | 0 |
| CGS | 50 | 50 | 23 | 12 | 1 |
| ZD | 50 | 50 | 0 | 0 | 0 |
| ZD+CGS | 50 | 50 | 0 | 0 | 1 |
| XE | 0 | 0 | 23 | 12 | 0 |
| XE+CGS | 0 | 0 | 23 | 12 | 1 |
| FSK | 50 | 50 | 23 | 12 | 1 |
| FSK+Lin | 50 | 20 | 23 | 12 | 1 |

### Model Reduction

We simplify the analysis of the system by exploiting the time scale separation in the model. Specifically, *V* is fast, *h* and *s* are slow, and *r* and *w* change with a super-slow rate. To analyze the fast subsystem, we treat *w* and *r* as model parameters and set them to their average values,

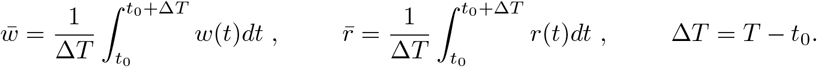

The simulation time interval (*T*) is set to 450 ms, while the initial transient (*t*_0_) is 350 ms. In order to be able to use the same analysis of the 3D fast subsystem both in the absence and presence of cAMP, we transform the gating variables *r* and *w* of the cAMP-sensitive M and HCN channels as follows. We introduce the scaling factors *G*_*w*_ and *G*_*r*_ and set

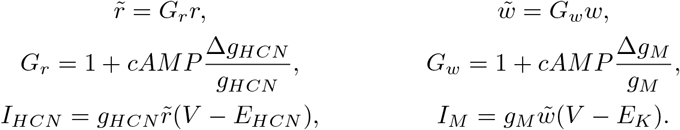

The same scaling operations are applied to map the 5D trajectories onto the 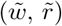 plane for comparison with 3D fast-subsystem BDs, in order to interpret the 5D model dynamics with the analyses made for the 3D model.

In conclusion, the reduced 3D model is

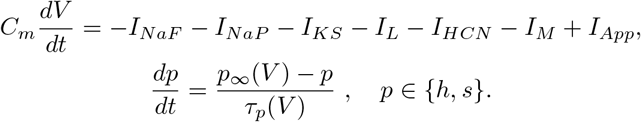

This system has itself multiple time scales, specifically, *V* is fast and *h* and *s* are slow. In this formulation, 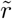 and 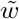 are parameters that assume values in the ranges [0, *G*_*r*_] and [0, *G*_*w*_], respectively.

### Multiple Time Scale Dynamical System

As noted above, the 3D fast subsystem is itself a slow-fast system with two slow and one fast variable. It has the standard structure [8, 25]

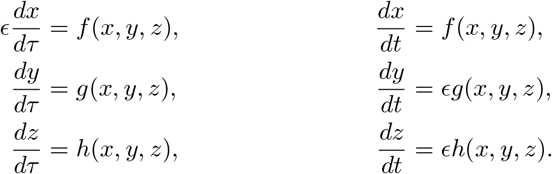

where 0 *< ϵ* ≪ 1 so *x* is the fast, and *y* and *z* are the slow. *t* and *τ* = *ϵt* are the fast and slow time scales, respectively. The associated solutions of the ODEs systems are known as slow and fast flows, respectively. By taking the limit *ϵ* → 0 is obtained,

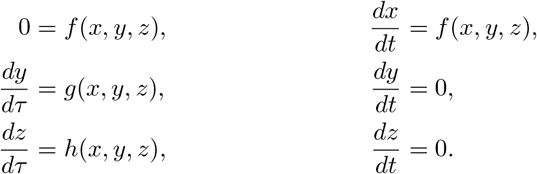

These are called reduced and layer problems, respectively. In the former, the system evolves according to the slow flow subject to the algebraic constraint 0 = *f* (*x, y, z*), while in the latter, the trajectory is governed by the fast dynamics. The critical manifold *𝒞*_0_ is the set of all points (*x, y, z*) in the phase space where *f* (*x, y, z*) = 0. This set represents the equilibrium points of the layer problem. The points in *𝒞*_0_ are attracting (repelling) if *∂*_*x*_*f* is negative (positive). Points with *∂*_*x*_*f* = 0 are said to be non-hyperbolic. The manifold *𝒞*_0_ is split into the attracting and repelling submanifolds, consisting of the attracting/repelling points, denoted *𝒮*_*a*_ and *𝒮*_*r*_, respectively. Generally, a set of non-hyperbolic points known as fold curves *ℒ* delimits these submanifolds,

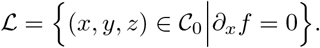

For positive but small *ϵ, 𝒞*_0_ perturbs to a slow manifold *𝒞*_*ϵ*_, which is *𝒪* (*ϵ*) close to _𝒞0_ [29]. Similarly, 𝒮_*a*_ and 𝒮_*r*_ perturb to 𝒮_*a,ϵ*_ and 𝒮_*r,ϵ*_. This relaxation perturbs the solutions of the reduced and the layer problems. Specifically, the trajectory is dominated by the slow (fast) dynamics when it is close to (away from) 𝒞_0_. A method to understand the dynamics near the fold, where the system becomes singular, consists in deriving the desingularized model. This model results from applying the total differentiation theorem, using the explicit expression of 𝒞_0_, and performing a time-rescaling known as desingularization, which yields [8, 25],

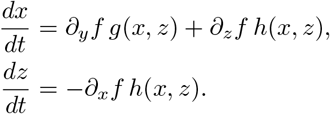

where 𝒞_0_ is expressed (locally) as *y* = *γ*(*x, z*). Equilibria of the desingularized system that satify *g*(*x, z*) = *h*(*x, z*) = 0 are also equilibria of the non-desingularized system and are called ordinary singularities. Instead, folded singularities satisfy *∂*_*x*_*f* = 0 and *∂*_*y*_*fg*(*x, z*) + *∂*_*z*_*fh*(*x, z*) = 0. The first condition imposes that the point lies on ℒ. A folded singularity is classified as a folded node (FN), folded saddle, or folded foci according to the eigenvalues of the Jacobian of the desingularized system. It is well known that FNs are associated with canard-induced SAOs, which can lead to MMOs with an appropriate return mechanism [30].

For the model analyzed here, the critical manifold can be written

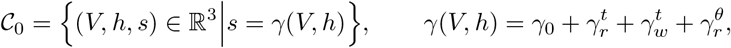

where the last expression presents a decomposition to understand better how variation in 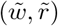 moves *𝒞*_0_. The terms are

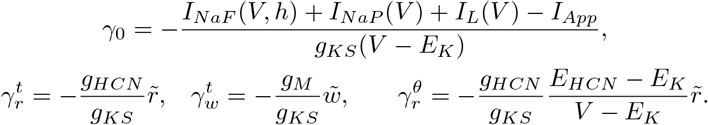

Here *γ*_0_ defines the basic structure of *𝒞*_0_ when both channels are inhibited. Instead, 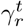 and 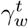 are shifting factors, representing how activation of HCN and M channels move *𝒞*_0_. Finally, 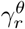applies a shift inversely proportional to *V*, but still proportional to 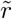. To split *𝒞*_0_ into its submanifolds, the fold *ℒ* is computed,

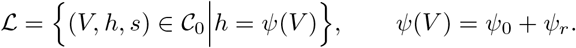

Again, the definition decomposes the fold line to evaluate how modification of HCN and M channel activation move *ℒ*,

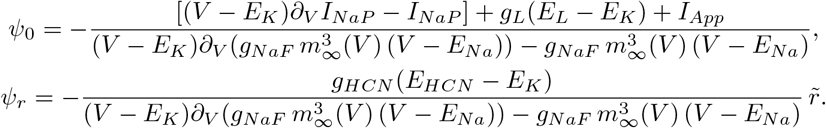

We see that the HCN gating variable 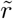 translates the fold line, whereas the *h* component of *ℒ* is M channel independent. However, because by construction the fold is constrained to live on 𝒞_0_, the *s* component of ℒ does depend on 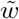 (as well as on 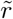) through *γ*, which considers the shift caused by the M channel activity.

Finally, the desingularized 3D model is

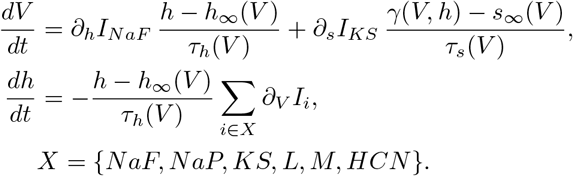

This system has two types of singularities. The ordinary singularities are obtained as the numerical solutions of *γ*(*V, h*_∞_(*V*)) = *s*_∞_(*V*). These points correspond to the equilibriums of the non-desingularized 3D model. Instead, the folded singularities are obtained by considering those points belonging to *ℒ* where

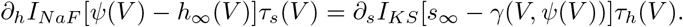

Solving this equation numerically provides the *V* coordinates of those points in ℒ classifiable as folded singularities. Numerically, we found that the desingularized system possesses a folded node singularity.

## Notes

### Competing Interest Statement

The authors have declared no competing interest.

